# Intrinsic multiplication rate variation of Plasmodium falciparum in clinical isolates prior to elimination in Malaysia

**DOI:** 10.1101/2024.12.10.627711

**Authors:** Lindsay B. Stewart, Elena Lantero Escolar, James Philpott, Janet Cox-Singh, Balbir Singh, David J. Conway

## Abstract

Replication rates and virulence of pathogens are hypothesised to evolve in response to varying intensity of transmission and competition among genotypes. Under exponential growth conditions in culture, clinical isolates of the malaria parasite *Plasmodium falciparum* have variable intrinsic multiplication rates, but comparisons of samples from different areas are needed. To analyse parasites from an area of low endemicity, Malaysian clinical isolates cryopreserved prior to malaria elimination were studied. The mean and range of *P. falciparum* multiplication rates in Malaysian isolates was no less than that seen among isolates from more highly endemic populations in Africa, which does not support a hypothesis of adaptation to prevailing levels of infection endemicity. Moreover, the distribution of multiplication rates was similar between isolates with single parasite genotypes and those containing multiple genotypes, which does not support a hypothesis of facultative adjustment to competing parasites. Based on clinical isolates, the findings indicate that parasites may not evolve lower multiplication rates under conditions of reduced transmission, and that the virulence potential is likely to be undiminished in pre-elimination settings. This encourages efforts to eliminate endemic infection completely, as has been achieved at the national level in Malaysia.

## 1. Introduction

It has been considered that pathogens may evolve high multiplication rates under conditions of intense transmission where superinfection is common, due to selection from within-host competition among different genotypes (Mackinnon and Read, 2004; Acevedo et al., 2019). Determining if this pertains to malaria has relevance to understanding and controlling the disease, caused by asexually replicating blood-stage parasites leading to high infection loads (Dondorp et al., 2005; Georgiadou et al., 2019). Most malaria cases globally are now caused by *Plasmodium falciparum* within Africa, although this parasite species also continues to be endemic in parts of Asia and South America, while increasing numbers of countries have become malaria-free (WHO, 2023). It is not known whether local parasite populations might evolve altered multiplication rates during the long process of malaria control required before elimination from any area.

Modelling of parasite and host parameters in studies of clinical malaria cases has suggested significant natural parasite variation in multiplication potential (Georgiadou et al., 2019), although intrinsic rates cannot be measured directly in such cases. Data from experimentally induced malaria infections have indicated variation in multiplication rates among *P. falciparum* strains, although protocol differences cause uncertainty in some comparisons (Simpson et al., 2002; Dietz et al., 2006; Friedman-Klabanoff et al., 2019; Laurens et al., 2019). Parasite strains maintained for a long time in laboratories often contain genomic alterations that do not occur in nature (Claessens et al., 2017), so comparisons of parasites in culture should ideally be performed within a short time of isolation (Brown and Guler, 2020). Assays of parasite multiplication in the first ex vivo cycle have indicated significant variation that may be associated with severity of disease, although such estimations are subject to variation in the viability of parasites in clinical samples (Chotivanich et al., 2000; Deans et al., 2006; Ribacke et al., 2013). After establishment of diverse clinical isolates in continuous culture, more controlled measurements under exponential growth conditions are possible, which have shown a range of *P. falciparum* multiplication rates from approximately 2-fold to 8-fold per 48 hours (Murray et al., 2017; Stewart et al., 2020).

Significant progress has been made towards elimination of *P. falciparum* infection in much of Southeast Asia. Since 2018, Malaysia has been free from any local transmission of *P. falciparum* or other human malaria parasites that used to be endemic, so that the only risk remains from imported infections or zoonotic malaria caused by different parasite species with macaque reservoir hosts (Yap et al., 2021; WHO, 2023). Significant reduction of endemic malaria in Malaysia occurred from the 1950s onwards, initially under the original WHO policy of Malaria Eradication which had an impact throughout the country, including Sabah state in which the annual number of cases was reduced from ∼250,000 in 1951 to ∼10,000 in 1969 (Rahman, 1982). After this period, malaria eradication was no longer considered achievable, but malaria became notifiable while it remained continuously endemic, with incident cases affecting less than 3% of the population in Sabah by the early 1990s. Subsequent intensification of malaria control led to further reduction of incidence (William et al., 2013), until the last indigenous case of *P. falciparum* or any non-zoonotic malaria in Malaysia was detected in 2017 (WHO, 2023).

It remains unknown whether reductions in malaria have predictable effects on parasite evolution. By the turn of the millennium, the *P. falciparum* population genetic structure in different areas of Malaysia including Sabah showed evidence of local fragmentation, and most *P. falciparum* infections contained single genotypes (Anthony et al., 2005). This contrasts with the situation in most of Africa where the parasite remains more highly endemic, and where most infections contain multiple genotypes due to superinfection as well as co-transmission of diverse genotypes (Zhu et al., 2019; Nkhoma et al., 2020; Wong et al., 2022; MalariaGen et al., 2023). To investigate parasite multiplication rate variation here, *P. falciparum* isolates were cultured from cryopreserved blood samples archived from cases sampled in Sabah in the late 1990s when the infection remained endemic. Exponential multiplication rates of the Malaysian parasites assayed in culture did not differ significantly between single and multiple-genotype isolates, and the overall distribution was similar to that seen for *P. falciparum* isolate samples from more highly endemic populations. These results do not indicate that parasite multiplication rates are adjusted in response to varying levels of endemicity or to the immediate presence of competing parasites.

## 2. Methods

### 2.1. Parasite sampling from malaria patients

Blood samples were collected from Plasmodium falciparum malaria cases attending government health facilities at three sites in Malaysian Borneo between December 1996 and December 1997 (Lahad Datu Hospital, Kunak Health Center, and Tawau General Hospital, in Southern Sabah) (Figure 1). Patients with *P. falciparum* infections were invited to donate 3ml of peripheral blood to contribute to understanding of variation in parasite genotypes and phenotypes including drug susceptibility (Cox-Singh et al., 2001). Recruitment of patients into the original study did not select by age or sex, but broadly reflected the cases presenting to each hospital including many adults. Most of the volume of each blood sample was collected into EDTA blood collection tubes, with aliquots being cryopreserved in liquid nitrogen as archival samples for laboratory culture, and patient data were since decoupled from these so that they were retained as a representative population sample of parasites. Cryopreserved blood samples for this study were maintained in liquid nitrogen in Malaysia until 2021, and then sent on dry ice to the London School of Hygiene and Tropical Medicine (LSHTM) for storage in liquid nitrogen until thawing for parasite culture. This process was approved by the Medical Research Ethics Committee of Universiti Malaysia Sarawak and the Ethics Committee of the London School of Hygiene and Tropical Medicine, in accordance with the International Conference of Harmonization Good Clinical Practice Guidelines.

**Fig 1.**
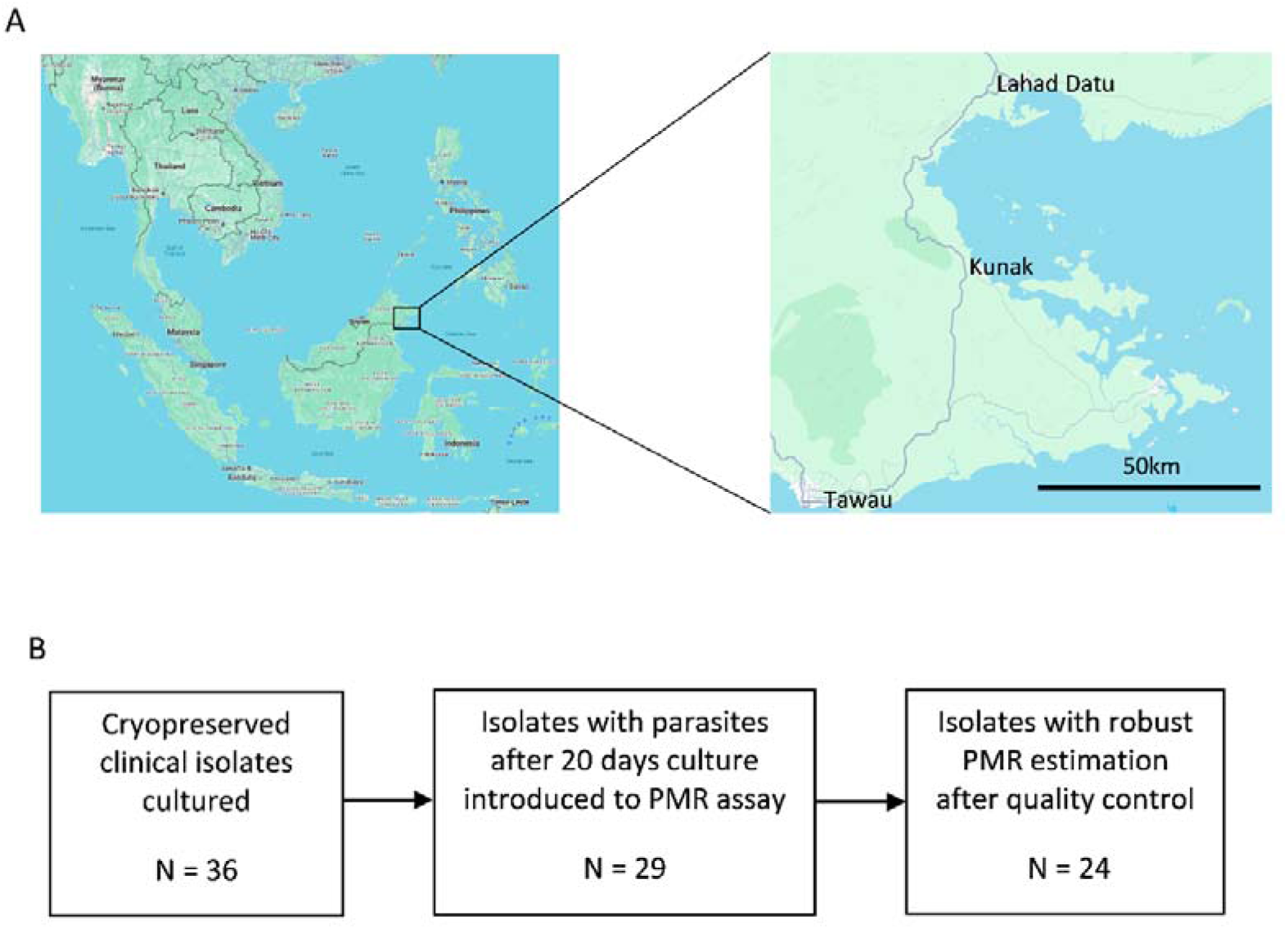
Sampling sites in Malaysia and laboratory processing of archived *P. falciparum* clinical samples. A. Map showing location of three hospital sites in the southeastern area of Sabah state in Borneo from which samples were cryopreserved (Kunak and Lahad Datu in 1996 and Tawau in 1997). Graphical map information from map data@2024 Google. B. Out of 36 isolates thawed and cultured (26 from Tawau, five from Kunak, and five from Lahad Datu), 29 had sufficient parasites in culture after 20 days to initiate 6-day assays of exponential multiplication rates, and parasite multiplication rates (PMR) per 48 hours were successfully derived for 24 of these isolates (18 from Tawau, four from Kunak, and two from Lahad Datu).

### 2.2. Parasite culture

Blood samples archived from each of 36 Malaysian *P. falciparum* malaria cases were thawed from glycerolyte cryopreservation and *P. falciparum* parasites were cultured continuously at 37 C, using standard methods. Briefly, a 12% NaCl solution was added dropwise to each cryopreserved sample while shaking the tube gently, adding up to half of the original sample volume, left to stand for 5 mins, following which 10 times the original volume of 1.6% NaCl was added dropwise to the sample while shaking the tube gently. After centrifugation for 5 min at 500 g, the cells were resuspended in RPMI 1640 medium containing 0.5% Albumax™ II (Thermo Fisher Scientific, Paisley, United Kingdom) and centrifuged again, following which the erythrocyte pellet was resuspended and combined with fresh erythrocytes from an anonymous donor of blood group O for culture at 3% haematocrit in RPMI 1640 medium supplemented with 0.5% Albumax II, under an atmosphere of 5% O_2_, 5% CO_2_, and 90% N_2_, with orbital shaking of flasks at 50 revolutions per minute. Fresh erythrocytes from anonymous donors of blood group O were added when cultures were diluted 1:5 at least once every week, so original patient erythrocytes would have virtually disappeared from cultures after a few weeks due to replacement caused by dilution. Isolates having parasites sustained continuously in culture, with at least 0.2% erythrocytes being infected after 20 days, were introduced into multiplication rate assays under experimental growth conditions as described below.

### 2.3. Parasite multiplication rate assays

Exponential multiplication rate assays were performed using a method previously described (Murray et al., 2017). Prior to each of the assays being initiated after at least 20 days of continuous culture, fresh blood was collected into EDTA tubes from three anonymous donors of blood group O, and erythrocytes were stored at 4 C for no more than 2 days and washed three times before use. Each of the assays for each isolate was performed in triplicate, with erythrocytes from each of the three different donors in separate flasks. Asynchronous parasite cultures were diluted to approximately 0.02% parasitaemia within each flask at the start of each assay which was conducted over six days. Every 48 hours (day 0, 2, 4, and 6) 200µl of suspended culture was taken for DNA extraction and qPCR, and culture media were renewed.

Following extraction of DNA from each of the day 0, 2, 4, and 6 timepoints in each of the assay replicates, qPCR to measure numbers of parasite genome copies was performed using a previously described protocol targeting a highly conserved locus in the *P. falciparum* genome (the *Pfs*25 gene) (Murray et al., 2017). Analysis of the parasite genome numbers at days 0, 2, 4 and 6 of each assay was performed by qPCR, and quality control was performed to exclude any assays with less than 100 parasite genome copy numbers measured at the end. Quality control also removed any points where a measurement was either lower or more than 20-fold higher than that from the same well two days earlier in the assay, and any outlying points among the biological triplicates that had a greater than two-fold difference from the other replicates on the same day. Assays were retained in the final analysis if there were duplicate or triplicate biological replicate measurements remaining after the quality control steps, and if they showed a coefficient of determination of *r*^2^ > 0.90 for the multiplication rate estimates using all remaining data points. For each assay, an overall parasite multiplication rate (defined as per 48-hour typical replicative cycle time) was calculated, with 95% confidence intervals using a standard linear model with GraphPad PRISM. The long-term laboratory adapted *P. falciparum* clone 3D7 was assayed in parallel as a control in all assays, consistently showing a multiplication rate of approximately 8.0 fold per 48 hours as described previously (Murray et al., 2017).

### 2.4. Parasite genotyping

To test for the presence of single or multiple *P. falciparum* genotypes within each of the isolates, genotyping of highly polymorphic repeat sequence regions within the *msp1* and *msp2* gene loci was performed on extracted genomic DNA from cultures sampled shortly before setting up the exponential multiplication rate assays. A nested PCR protocol incorporated msp1 block 2 and *msp2* dimorphic region allelic family-specific primers, and alleles were discriminated by scoring of PCR product sizes and allelic family types on agarose gels (2% agarose for *msp2* allelic products, and 3% agarose for msp1 allelic products) using a method previously described (Snounou et al., 1999). This method gives usefully sensitive and reproducible detection of mixed-genotype isolates, as previously compared with other convenient genotyping methods (Farnert et al., 2001). These two loci are known to have many different alleles within Southeast Asia as well as in Africa (Ferreira and Hartl, 2007; Aspeling-Jones and Conway, 2018), enabling discrimination of most parasite genotypes. Due to the sensitive nested PCR and high level of allelic discrimination, the method can also detect mixed genotype infections not clearly apparent by whole-genome sequencing of isolates as some genotypes are present in low minority proportions (Auburn et al., 2012).

## 3. Results

### 3.1. Culture establishment and assay of *P. falciparum* isolates from archived Malaysian samples

Thirty-six archived Malaysian *P. falciparum* clinical blood samples that had been cryopreserved in 1996-1997 were thawed and parasites introduced into culture, involving dilution and replacement into new erythrocytes at culture initiation and periodically to support continuous growth of the parasites. Twenty-nine (81%) of these isolates had parasites continuously growing after 20 days of continuous culture, at a density of at least 0.2% erythrocytes infected, which was sufficient to allow dilution to very low parasitaemia for assay of multiplication rates (Figure 1). Each of these isolates was tested in a 6-day exponential multiplication rate assay at starting parasitaemia of 0.02% in triplicate flasks with erythrocytes from three different donors. After filtering for assay quality control, the data for 24 of the isolates passed the criteria for estimation of exponential parasite multiplication rates (Figure 2 and Supplementary Table S1).

**Fig 2.**
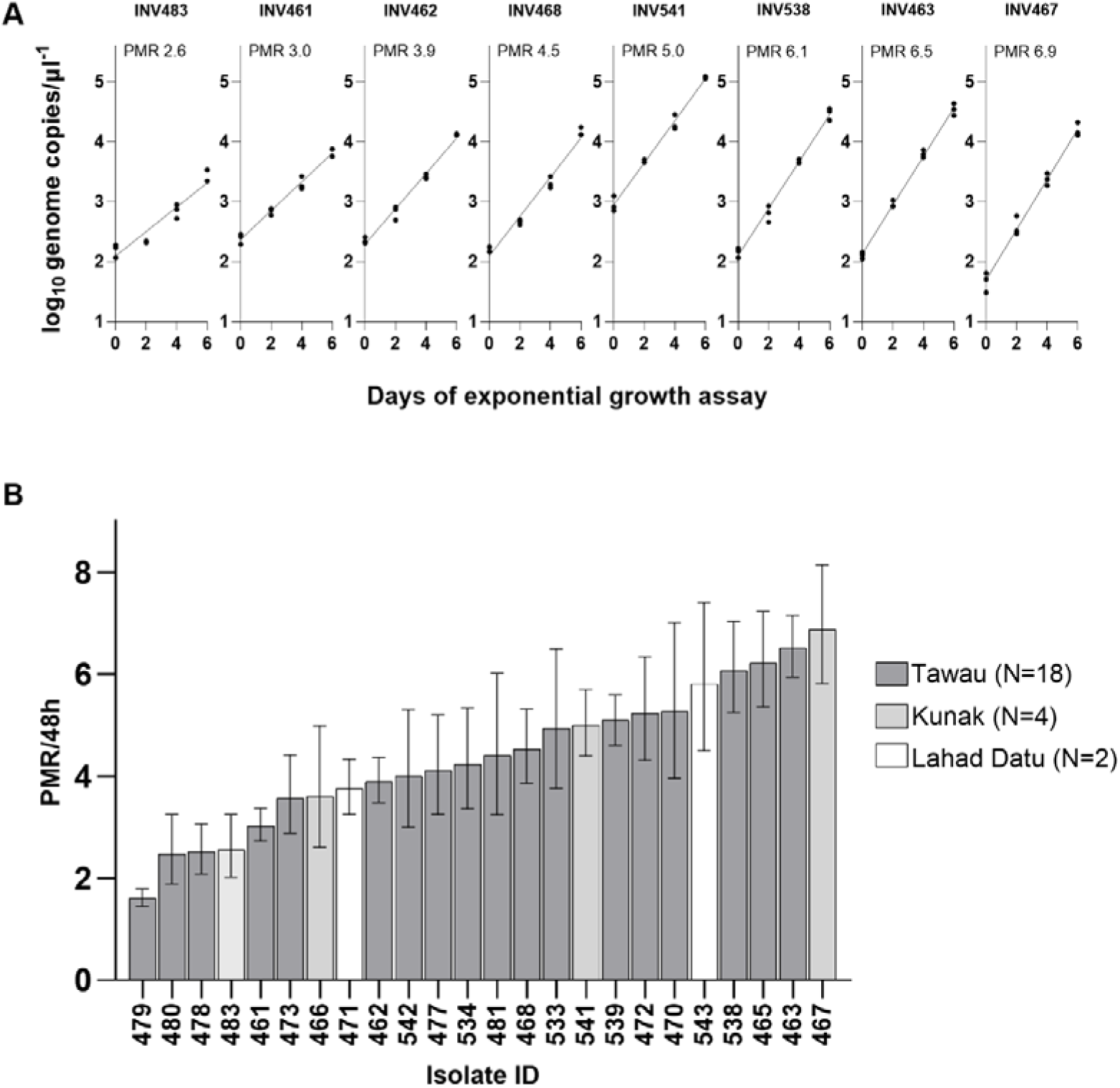
Variation in *P. falciparum* multiplication rates among cultured clinical isolates from Malaysia. A. Exponential growth assay data illustrated for 8 isolates (plots for all 24 isolates are shown in Supplementary Figure S1), tested in a 6-day assay conducted in triplicate with erythrocytes from three different donors. Plots show parasite genome copies (Log_10_ scale) per microlitre of DNA extracted from each culture timepoint). Assays were performed after each isolate had been in continuous culture for least 20 and no more than 23 days (Supplementary Table S1). Estimates of parasite multiplication rates (PMR) per 48 hours for each isolate are derived by logistic regression. B. Bar chart of multiplication rates and 95% confidence intervals of estimates for all 24 individual isolates ranging from the lowest (PMR = 1.6) to the highest (PMR = 6.9). Values are given in Supplementary Table S1.

### 3.2. Multiplication rate variation among Malaysian *P. falciparum* isolates

Exponential parasite multiplication rates across the 24 Malaysian isolates ranged from 1.6-fold to 6.9-fold (per 48-hours corresponding to a typical asexual cycle time) (Figures 2 and 3). For each isolate, there was generally high consistency in the parasite genome copy numbers measured in parallel replicate cultures with different erythrocyte donors (Figure 2), so the multiplication rate estimates had tight confidence intervals (Figure 3 and Supplementary Table S1). Across all isolates there was an overall mean of 4.4-fold multiplication per 48 hours (Figure 3). Most of the isolates were from Tawau hospital, but the isolates from the other sites did not show a trend towards having higher or lower rates (Figure 3), indicating that the variation is not due to differences between hospitals sampled (all were within 100 km of each other as shown in Figure 1).

**Fig 3.**
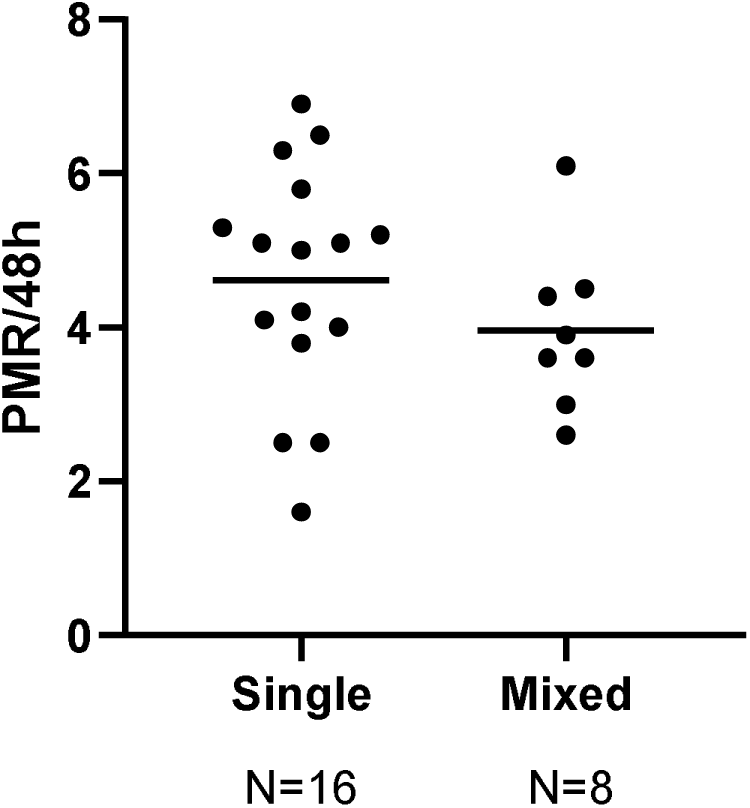
Comparison of multiplication rates in Malaysian *P. falciparum* isolates with single and multiple genotypes detected (Mann-Whitney P = 0.21). Detection of multiple genotypes was based on discrimination of alleles of msp1 and msp2 after PCR amplification. Numbers of alleles detected in each isolate for each locus are shown in Supplementary Table S2.

### 3.3. Comparison between isolates with single and multiple genotypes

Nine (36%) of the 24 isolates assayed contained multiple *P. falciparum* genotypes, as determined by allelic discrimination at the highly polymorphic *msp1* and *msp2* loci, while the remaining sixteen isolates each had a single parasite genotype detected (Figure 4 and Supplementary Table S2). There was no significant difference in parasite multiplication rates in comparison between the multiple genotype isolates (mean = 3.9-fold) and single genotype isolates (mean = 4.6-fold) (Figure 4).

**Fig 4.**
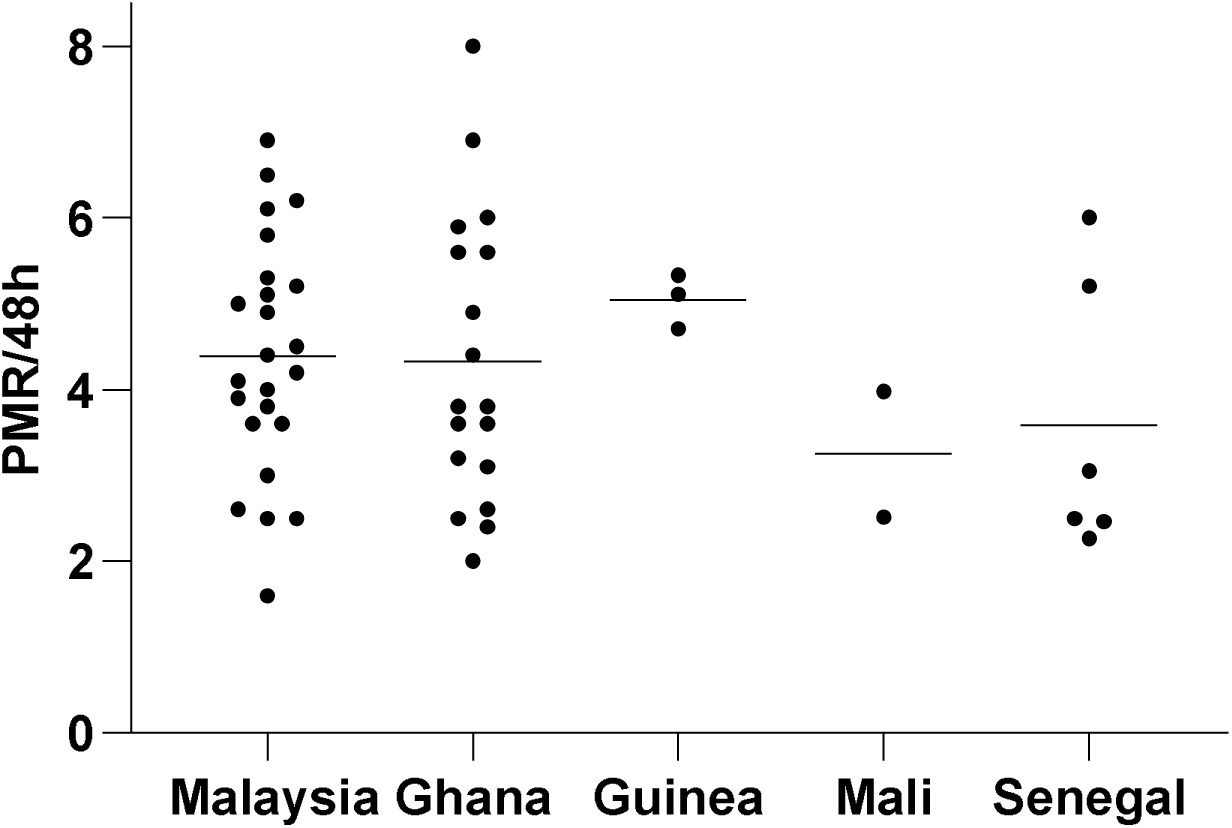
Comparison of multiplication rates of Malaysian isolates (Mean PMR = 4.4, N=24) with previous data for isolates from Ghana (Mean PMR = 4.3, N=18), Guinea (Mean PMR = 5.1, N=3), Mali (Mean PMR = 3.2, N=2) and Senegal (Mean PMR = 3.6, N = 6) tested under similar assay conditions. Horizontal lines show mean values for each country. The isolates from Ghana were assayed after 25 days of culture (Stewart et al., 2020), and those from elsewhere in West Africa were assayed after a minimum of 16 days and a mean of 23 days in culture (Murray et al., 2017). There was no significant difference in the distributions of PMR values for Malaysia compared with Ghana (Mann-Whitney P = 0.6) or compared with the rest of the West African isolates combined (Mann-Whitney P = 0.4).

A diagnostic assessment of parasitaemia in the peripheral blood samples of patients had been recorded using a categorical grading system based on numbers of parasites observed microscopically per high power field in the health facilities (Supplementary Table S2). Among the 22 isolates for which such records were available, there was a weak and non-significant correlation between parasite density in the patients and the multiplication rates measured in culture (Spearman’s rho = 0.12, P = 0.6)(Supplementary Figure S2).

### 3.4. Comparison of Malaysian isolates with those from more highly-endemic populations

The overall distribution of multiplication rates of the Malaysian isolates was compared with data on isolates sampled from West African populations between 2011 and 2013, that were previously determined using the same assay methods in the same laboratory (Murray et al., 2017; Stewart et al., 2020) (Figure 5). The values for Malaysian isolates showed a similar distribution to that seen for isolates sampled from a highly-endemic area of northern Ghana (N = 18, mean of 4.3-fold, Mann-Whitney P = 0.6) (Figure 5). The values for the Malaysian isolates were also similar to those from a more diverse panel of clinical isolates from three other countries in West Africa (N = 11, mean of 4.2-fold, Mann-Whitney P = 0.4) (Figure 5). Both of these comparisons show that Malaysian parasites did not have intrinsically lower multiplication rates than *P. falciparum* isolates from the most highly endemic region in the world.

## 4. Discussion

Understanding parasite multiplication rate variation is advanced here by analyses of samples collected in Malaysia when *P. falciparum* infection was endemic. Together with data on *P. falciparum* isolates from other populations, the results do not support a simple hypothesis suggesting directional trait adaptation, nor of facultative adjustment in the presence of competing parasites. It is clear that the mean and range of Malaysian parasite multiplication rates was similar to that of parasites from more highly endemic populations. Moreover, the variation in parasite multiplication rates among the Malaysian parasites did not differ for single genotype and multiple genotype isolates.

It is important to note that all of the archived Malaysian samples available were clinical isolates from patients, whereas it remains possible that parasites in asymptomatic infections in the community may have had lower multiplication rates. It is more difficult to study phenotypes of parasites in isolates of community infections as they are usually at very low levels in the blood (Imwong et al., 2016) and may be slower replicating (Stewart et al., 2024), but such infections are important to consider as malaria control proceeds (Bjorkman and Morris, 2020; Whittaker et al., 2021). The isolates from Malaysia were archived approximately 15 years before the previously analysed isolates from Africa, whereas sampling parasites nearer to the time of eventual local elimination might possibly show differences due to continuing parasite evolution, although small effective population sizes would reduce the likelihood of adaptive changes. Where feasible, it may be worth sampling asymptomatic infections to study parasite phenotypes elsewhere prior to elimination.

Although parasite multiplication rates appear to have similar distributions in different endemic populations, the variation among isolates within each local population may potentially have adaptive and clinical significance. A significant positive correlation between multiplication rates in culture and peripheral blood parasitaemia levels at time of sampling was seen among isolates from clinical cases in Ghana (Stewart et al., 2020), and studies of first cycle *ex vivo* replication in Thailand and Uganda had shown higher rates in parasites from severe malaria compared to mild malaria isolates (Chotivanich et al., 2000; Ribacke et al., 2013). In the present study on archived Malaysian samples, the test for a correlation between multiplication rates and patient parasitaemia is limited by the available semi-quantitative estimates of the latter in health facilities at the time of sampling. Prospective studies where malaria is still endemic will benefit from more intensive protocols to estimate densities in peripheral blood, or biomarker methods to estimate total parasite biomass (Dondorp et al., 2005; Georgiadou et al., 2019), which will help explore associations with intrinsic parasite multiplication rates.

The substantial observed variation in multiplication rates of clinical isolates is not primarily explainable by parasites undergoing sexual commitment. Parasite sexual commitment was not tested in this study, but previous studies have consistently shown that normally only a minority of *P. falciparum* parasites commit to sexual development, less than 20% per cycle (Stewart et al., 2022). A study of gene expression has suggested parasites may increase commitment to developing sexual stages to infect mosquitoes when overall transmission rates are lower (Abdi et al., 2023), which could marginally decrease asexual multiplication rates. The present study does not exclude the possibility of such an effect, as it would be too slight to be detected here.

Parasites in clinical isolates here were compared after only a few weeks in culture, giving sufficient time for erythrocytes from patients to have been diluted out by the time of assay, but not time for parasite mutants to become common (Claessens et al., 2017; Stewart et al., 2020; Claessens et al., 2023). However, any assessment of multiplication rates performed in culture might not fully reflect the multiplication potential of parasites in the in vivo environment within patients. A general increase in replication rates over several months of continuous culture was previously seen for Ghanaian isolates, which might either reflect phenotype plasticity due to epigenetic regulation or effects of genomic changes, the relative importance of which remains to be determined (Stewart et al., 2020; Claessens et al., 2023). It is also possible that other variables may determine differences between parasite multiplication rates under non-exponential growth conditions, when there would be significant density-dependent effects (Rovira-Graells et al., 2016; Gnangnon et al., 2021). For example, significant variation in relative growth rates of parasite isolates from the Thai-Myanmar border was seen when a small panel was tested in pairwise combinations under competitive conditions (Tirrell et al., 2019). More comparisons of parasite relative growth rates under different culture conditions may reveal variation that could reflect adaptive changes.

At the cellular level, it will be important to discover which phases of the replication and growth cycle contribute most to the variation of *P. falciparum* multiplication rates. An initial analysis of different Ghanaian isolates indicated that the number of merozoites within mature schizonts varies only slightly and was not a major determinant of the variation in multiplication rates (Stewart et al., 2020). Other variables to be investigated in terms of their relative contribution include cell cycle duration (Reilly Ayala et al., 2010; Ganter et al., 2017; Gnangnon et al., 2021) and invasion efficiency of merozoites (Gnangnon et al., 2021), including the use of alternative erythrocyte receptors for invasion which shows some variation among populations (Bowyer et al., 2015). Analysis of parasites from different populations is advisable to ensure that mechanisms characterised *in vitro* reflect the basis of natural phenotypic variation.

## Acknowledgements

We are grateful to Khamisah Binti Abdul Kadir for managing the cryopreserved collection of isolates at UNIMAS, and arranging the sample documentation and information for sample shipment. The study was supported by funding from the UK Medical Research Council (Project grant MR/S009760/1).

## CRediT authorship contribution statement

Lindsay B. Stewart: Conceptualization, Data curation, Formal analysis, Investigation, Methodology, Project administration, Resources, Validation, Writing – original draft, Writing – review and editing.

Elena Lantero-Escolar: Formal analysis, Investigation, Methodology, Validation, Writing – review and editing.

James Philpott: Investigation, Writing – review and editing.

Janet Cox-Singh: Investigation, Writing – review and editing.

Balbir Singh: Investigation, Writing – review and editing.

David J. Conway: Conceptualization, Data curation, Formal analysis, Funding acquisition, Investigation, Methodology, Project administration, Resources, Supervision, Validation, Writing – original draft, Writing – review and editing.

## References

Abdi, A.I., Achcar, F., Sollelis, L., Silva-Filho, J.L., Mwikali, K., Muthui, M., Mwangi, S., Kimingi, H.W., Orindi, B., Andisi Kivisi, C., Alkema, M., Chandrasekar, A., Bull, P.C., Bejon, P., Modrzynska, K., Bousema, T., Marti, M., 2023. Plasmodium falciparum adapts its investment into replication versus transmission according to the host environment. Elife 12. doi: 10.7554/eLife.85140

Acevedo, M.A., Dillemuth, F.P., Flick, A.J., Faldyn, M.J., Elderd, B.D., 2019. Virulence-driven trade-offs in disease transmission: A meta-analysis. Evolution 73, 636–647. doi: 10.1111/evo.13692

Anthony, T.G., Conway, D.J., Cox-Singh, J., Matusop, A., Ratnam, S., Shamsul, S., Singh, B., 2005. Fragmented population structure of Plasmodium falciparum in a region of declining endemicity. J Infect Dis 191, 1558–1564.

Aspeling-Jones, H., Conway, D.J., 2018. An expanded global inventory of allelic variation in the most extremely polymorphic region of Plasmodium falciparum merozoite surface protein 1 provided by short read sequence data. Malaria Journal 17, 345. doi: 10.1186/s12936-018-2475-2

Auburn, S., Campino, S., Miotto, O., Djimde, A.A., Zongo, I., Manske, M., Maslen, G., Mangano, V., Alcock, D., MacInnis, B., Rockett, K.A., Clark, T.G., Doumbo, O.K., Ouedraogo, J.B., Kwiatkowski, D.P., 2012. Characterization of within-host Plasmodium falciparum diversity using next-generation sequence data. PLoS One 7, e32891. doi: 10.1371/journal.pone.0032891

Bjorkman, A., Morris, U., 2020. Why asymptomatic Plasmodium falciparum infections are common in low-transmission settings. Trends Parasitol 36, 898–905. doi: 10.1016/j.pt.2020.07.008

Bowyer, P.W., Stewart, L.B., Aspeling-Jones, H., Mensah-Brown, H.E., Ahouidi, A.D., Amambua-Ngwa, A., Awandare, G.A., Conway, D.J., 2015. Variation in Plasmodium falciparum erythrocyte invasion phenotypes and merozoite ligand gene expression across different populations in areas of malaria endemicity. Infect Immun 83, 2575–2582. doi: 10.1128/IAI.03009-14

Brown, A.C., Guler, J.L., 2020. From circulation to cultivation: Plasmodium in vivo versus in vitro. Trends Parasitol 36, 914–926. doi: 10.1016/j.pt.2020.08.008

Chotivanich, K., Udomsangpetch, R., Simpson, J.A., Newton, P., Pukrittayakamee, S., Looareesuwan, S., White, N.J., 2000. Parasite multiplication potential and the severity of falciparum malaria. J Infect Dis 181, 1206–1209.

Claessens, A., Affara, M., Assefa, S.A., Kwiatkowski, D.P., Conway, D.J., 2017. Culture adaptation of malaria parasites selects for convergent loss-of-function mutants. Sci Rep 7, 41303. doi: 10.1038/srep41303

Claessens, A., Stewart, L.B., Drury, E., Ahouidi, A.D., Amambua-Ngwa, A., Diakite, M., Kwiatkowski, D.P., Awandare, G.A., Conway, D.J., 2023. Genomic variation during culture adaptation of genetically complex Plasmodium falciparum clinical isolates. Microb Genom 9, e001009. doi: 10.1099/mgen.0.001009

Cox-Singh, J., Zakaria, R., Abdullah, M.S., Rahman, H.A., Nagappan, S., Singh, B., 2001. Short report: differences in dihydrofolate reductase but not dihydropteroate synthase alleles in Plasmodium falciparum isolates from geographically distinct areas in Malaysia. American Journal of Tropical Medicine and Hygiene 64, 28–31.

Deans, A.M., Lyke, K.E., Thera, M.A., Plowe, C.V., Kone, A., Doumbo, O.K., Kai, O., Marsh, K., Mackinnon, M.J., Raza, A., Rowe, J.A., 2006. Low multiplication rates of African Plasmodium falciparum isolates and lack of association of multiplication rate and red blood cell selectivity with malaria virulence. Am J Trop Med Hyg 74, 554–563.

Dietz, K., Raddatz, G., Molineaux, L., 2006. Mathematical model of the first wave of Plasmodium falciparum asexual parasitemia in non-immune and vaccinated individuals. Am J Trop Med Hyg 75, 46–55.

Dondorp, A.M., Desakorn, V., Pongtavornpinyo, W., Sahassananda, D., Silamut, K., Chotivanich, K., Newton, P.N., Pitisuttithum, P., Smithyman, A.M., White, N.J., Day, N.P., 2005. Estimation of the total parasite biomass in acute falciparum malaria from plasma PfHRP2. PLoS Med 2, e204. doi:

Farnert, A., Arez, A.P., Babiker, H.A., Beck, H.P., Benito, A., Bjorkman, A., Bruce, M.C., Conway, D.J., Day, K.P., Henning, L., Mercereau-Puijalon, O., Ranford-Cartwright, L.C., Rubio, J.M., Snounou, G., Walliker, D., Zwetyenga, J., do Rosario, V.E., 2001. Genotyping of Plasmodium falciparum infections by PCR: a comparative multicentre study. Trans R Soc Trop Med Hyg 95, 225–232.

Ferreira, M.U., Hartl, D.L., 2007. Plasmodium falciparum: worldwide sequence diversity and evolution of the malaria vaccine candidate merozoite surface protein-2 (MSP-2). Exp Parasitol 115, 32–40.

Friedman-Klabanoff, D.J., Laurens, M.B., Berry, A.A., Travassos, M.A., Adams, M., Strauss, K.A., Shrestha, B., Levine, M.M., Edelman, R., Lyke, K.E., 2019. The Controlled Human Malaria Infection experience at the University of Maryland. Am J Trop Med Hyg 100, 556–565. doi: 10.4269/ajtmh.18-0476

Ganter, M., Goldberg, J.M., Dvorin, J.D., Paulo, J.A., King, J.G., Tripathi, A.K., Paul, A.S., Yang, J., Coppens, I., Jiang, R.H., Elsworth, B., Baker, D.A., Dinglasan, R.R., Gygi, S.P., Duraisingh, M.T., 2017. Plasmodium falciparum CRK4 directs continuous rounds of DNA replication during schizogony. Nat Microbiol 2, 17017. doi: 10.1038/nmicrobiol.2017.17

Georgiadou, A., Lee, H.J., Walther, M., van Beek, A.E., Fitriani, F., Wouters, D., Kuijpers, T.W., Nwakanma, D., D’Alessandro, U., Riley, E.M., Otto, T.D., Ghani, A., Levin, M., Coin, L.J., Conway, D.J., Bretscher, M.T., Cunnington, A.J., 2019. Modelling pathogen load dynamics to elucidate mechanistic determinants of host-Plasmodium falciparum interactions. Nat Microbiol 4, 1592–1602. doi: 10.1038/s41564-019-0474-x

Gnangnon, B., Duraisingh, M.T., Buckee, C.O., 2021. Deconstructing the parasite multiplication rate of Plasmodium falciparum. Trends Parasitol 37, 922–932. doi: 10.1016/j.pt.2021.05.001

Imwong, M., Stepniewska, K., Tripura, R., Peto, T.J., Lwin, K.M., Vihokhern, B., Wongsaen, K., von Seidlein, L., Dhorda, M., Snounou, G., Keereecharoen, L., Singhasivanon, P., Sirithiranont, P., Chalk, J., Nguon, C., Day, N.P., Nosten, F., Dondorp, A., White, N.J., 2016. Numerical distributions of parasite densities during asymptomatic malaria. J Infect Dis 213, 1322–1329. doi: 10.1093/infdis/jiv596

Laurens, M.B., Berry, A.A., Travassos, M.A., Strauss, K., Adams, M., Shrestha, B., Li, T., Eappen, A., Manoj, A., Abebe, Y., Murshedkar, T., Gunasekera, A., Richie, T.L., Lyke, K.E., Plowe, C.V., Kennedy, J.K., Potter, G.E., Deye, G.A., Sim, B.K.L., Hoffman, S.L., 2019. Dose-dependent Infectivity of aseptic, purified, cryopreserved Plasmodium falciparum 7G8 sporozoites in malaria-naive adults. J Infect Dis 220, 1962–1966. doi: 10.1093/infdis/jiz410

Mackinnon, M.J., Read, A.F., 2004. Virulence in malaria: an evolutionary viewpoint. Philos Trans R Soc Lond B Biol Sci 359, 965–986.

MalariaGen, Abdel Hamid, M.M., Abdelraheem, M.H., Acheampong, D.O., Ahouidi, A., Ali, M., Almagro-Garcia, J., Amambua-Ngwa, A., Amaratunga, C., Amenga-Etego, L., Andagalu, B., Anderson, T., Andrianaranjaka, V., Aniebo, I., Aninagyei, E., Ansah, F., Ansah, P.O., Apinjoh, T., Arnaldo, P., Ashley, E., Auburn, S., Awandare, G.A., Ba, H., Baraka, V., Barry, A., Bejon, P., Bertin, G.I., Boni, M.F., Borrmann, S., Bousema, T., Bouyou-Akotet, M., Branch, O., Bull, P.C., Cheah, H., Chindavongsa, K., Chookajorn, T., Chotivanich, K., Claessens, A., Conway, D.J., Corredor, V., Courtier, E., Craig, A., D’Alessandro, U., Dama, S., Day, N., Denis, B., Dhorda, M., Diakite, M., Djimde, A., Dolecek, C., Dondorp, A., Doumbia, S., Drakeley, C., Drury, E., Duffy, P., Echeverry, D.F., Egwang, T.G., Enosse, S.M.M., Erko, B., Fairhurst, R.M., Faiz, A., Fanello, C.A., Fleharty, M., Forbes, M., Fukuda, M., Gamboa, D., Ghansah, A., Golassa, L., Goncalves, S., Harrison, G.L.A., Healy, S.A., Hendry, J.A., Hernandez-Koutoucheva, A., Hien, T.T., Hill, C.A., Hombhanje, F., Hott, A., Htut, Y., Hussein, M., Imwong, M., Ishengoma, D., Jackson, S.A., Jacob, C.G., Jeans, J., Johnson, K.J., Kamaliddin, C., Kamau, E., Keatley, J., Kochakarn, T., Konate, D.S., Konate, A., Kone, A., Kwiatkowski, D.P., Kyaw, M.P., Kyle, D., Lawniczak, M., Lee, S.K., Lemnge, M., Lim, P., Lon, C., Loua, K.M., Mandara, C.I., Marfurt, J., Marsh, K., Maude, R.J., Mayxay, M., Maiga-Ascofare, O., Miotto, O., Mita, T., Mobegi, V., Mohamed, A.O., Mokuolu, O.A., Montgomery, J., Morang’a, C.M., Mueller, I., Murie, K., Newton, P.N., Ngo Duc, T., Nguyen, T., Nguyen, T.N., Nguyen Thi Kim, T., Nguyen Van, H., Noedl, H., Nosten, F., Noviyanti, R., Ntui, V.N., Nzila, A., Ochola-Oyier, L.I., Ocholla, H., Oduro, A., Omedo, I., Onyamboko, M.A., Ouedraogo, J.B., Oyebola, K., Oyibo, W.A., Pearson, R., Peshu, N., Phyo, A.P., Plowe, C.V., Price, R.N., Pukrittayakamee, S., Quang, H.H., Randrianarivelojosia, M., Rayner, J.C., Ringwald, P., Rosanas-Urgell, A., Rovira-Vallbona, E., Ruano-Rubio, V., Ruiz, L., Saunders, D., Shayo, A., Siba, P., Simpson, V.J., Sissoko, M.S., Smith, C., Su, X.Z., Sutherland, C., Takala-Harrison, S., Talman, A., Tavul, L., Thanh, N.V., Thathy, V., Thu, A.M., Toure, M., Tshefu, A., Verra, F., Vinetz, J., Wellems, T.E., Wendler, J., White, N.J., Whitton, G., Yavo, W., van der Pluijm, R.W., 2023. Pf7: an open dataset of Plasmodium falciparum genome variation in 20,000 worldwide samples. Wellcome Open Res 8, 22. doi: 10.12688/wellcomeopenres.18681.1

Murray, L., Stewart, L.B., Tarr, S.J., Ahouidi, A.D., Diakite, M., Amambua-Ngwa, A., Conway, D.J., 2017. Multiplication rate variation in the human malaria parasite Plasmodium falciparum. Sci Rep 7, 6436. doi: 10.1038/s41598-017-06295-9

Nkhoma, S.C., Trevino, S.G., Gorena, K.M., Nair, S., Khoswe, S., Jett, C., Garcia, R., Daniel, B., Dia, A., Terlouw, D.J., Ward, S.A., Anderson, T.J.C., Cheeseman, I.H., 2020. Co-transmission of related malaria parasite lineages shapes within-host parasite diversity. Cell Host Microbe 27, 93–103 e104. doi: 10.1016/j.chom.2019.12.001

Rahman, K.M., 1982. Epidemiology of malaria in Malaysia. Rev Infect Dis 4, 985–991. doi: 10.1093/clinids/4.5.985

Reilly Ayala, H.B., Wacker, M.A., Siwo, G., Ferdig, M.T., 2010. Quantitative trait loci mapping reveals candidate pathways regulating cell cycle duration in Plasmodium falciparum. BMC Genomics 11, 577. doi: 10.1186/1471-2164-11-577

Ribacke, U., Moll, K., Albrecht, L., Ahmed Ismail, H., Normark, J., Flaberg, E., Szekely, L., Hultenby, K., Persson, K.E., Egwang, T.G., Wahlgren, M., 2013. Improved in vitro culture of Plasmodium falciparum permits establishment of clinical isolates with preserved multiplication, invasion and rosetting phenotypes. PLoS One 8, e69781. doi: 10.1371/journal.pone.0069781

Rovira-Graells, N., Aguilera-Simon, S., Tinto-Font, E., Cortes, A., 2016. New assays to characterise growth-related phenotypes of Plasmodium falciparum reveal variation in density-dependent growth inhibition between parasite lines. PLoS One 11, e0165358. doi: 10.1371/journal.pone.0165358

Simpson, J.A., Aarons, L., Collins, W.E., Jeffery, G.M., White, N.J., 2002. Population dynamics of untreated Plasmodium falciparum malaria within the adult human host during the expansion phase of the infection. Parasitology 124, 247–263.

Snounou, G., Zhu, X., Siripoon, N., Jarra, W., Thaithong, S., Brown, K.N., Viriyakosol, S., 1999. Biased distribution of msp1 and msp2 allelic variants in Plasmodium falciparum populations in Thailand. Trans R Soc Trop Med Hyg 93, 369–374.

Stewart, L.B., Diaz-Ingelmo, O., Claessens, A., Abugri, J., Pearson, R.D., Goncalves, S., Drury, E., Kwiatkowski, D.P., Awandare, G.A., Conway, D.J., 2020. Intrinsic multiplication rate variation and plasticity of human blood stage malaria parasites. Commun Biol 3, 624. doi: 10.1038/s42003-020-01349-7

Stewart, L.B., Freville, A., Voss, T.S., Baker, D.A., Awandare, G.A., Conway, D.J., 2022. Plasmodium falciparum sexual commitment rate variation among clinical isolates and diverse laboratory-adapted lines. Microbiol Spectr, e0223422. doi: 10.1128/spectrum.02234-22

Stewart, L.B., Lantero Escolar, E., Philpott, J., Claessens, A., Amambua-Ngwa, A., Conway, D.J., 2024. Higher multiplication rates of Plasmodium falciparum in isolates from hospital cases compared with community infections. BioRxiv. doi: 10.1101/2024.05.02.592253

Tirrell, A.R., Vendrely, K.M., Checkley, L.A., Davis, S.Z., McDew-White, M., Cheeseman, I.H., Vaughan, A.M., Nosten, F.H., Anderson, T.J.C., Ferdig, M.T., 2019. Pairwise growth competitions identify relative fitness relationships among artemisinin resistant Plasmodium falciparum field isolates. Malar J 18, 295. doi: 10.1186/s12936-019-2934-4

Whittaker, C., Slater, H., Nash, R., Bousema, T., Drakeley, C., Ghani, A.C., Okell, L.C., 2021. Global patterns of submicroscopic Plasmodium falciparum malaria infection: insights from a systematic review and meta-analysis of population surveys. Lancet Microbe 2, e366–e374. doi: 10.1016/S2666-5247(21)00055-0

WHO, 2023. World Malaria Report 2023, Geneva.

William, T., Rahman, H.A., Jelip, J., Ibrahim, M.Y., Menon, J., Grigg, M.J., Yeo, T.W., Anstey, N.M., Barber, B.E., 2013. Increasing incidence of Plasmodium knowlesi malaria following control of *P. falciparum* and P. vivax Malaria in Sabah, Malaysia. PLoS Negl Trop Dis 7, e2026. 10.1371/journal.pntd.0002026

Wong, W., Volkman, S., Daniels, R., Schaffner, S., Sy, M., Ndiaye, Y.D., Badiane, A.S., Deme, A.B., Diallo, M.A., Gomis, J., Sy, N., Ndiaye, D., Wirth, D.F., Hartl, D.L., 2022. R (H): a genetic metric for measuring intrahost Plasmodium falciparum relatedness and distinguishing cotransmission from superinfection. PNAS Nexus 1, pgac187. doi: 10.1093/pnasnexus/pgac187

Yap, N.J., Hossain, H., Nada-Raja, T., Ngui, R., Muslim, A., Hoh, B.P., Khaw, L.T., Kadir, K.A., Simon Divis, P.C., Vythilingam, I., Singh, B., Lim, Y.A., 2021. Natural human infections with Plasmodium cynomolgi, P. inui, and 4 other simian malaria parasites, Malaysia. Emerg Infect Dis 27, 2187–2191. doi: 10.3201/eid2708.204502

Zhu, S.J., Hendry, J.A., Almagro-Garcia, J., Pearson, R.D., Amato, R., Miles, A., Weiss, D.J., Lucas, T.C., Nguyen, M., Gething, P.W., Kwiatkowski, D., McVean, G., Pf3k, P., 2019. The origins and relatedness structure of mixed infections vary with local prevalence of *P. falciparum* malaria. Elife 8, e40845. doi: 10.7554/eLife.40845

