## Supplementary for "Intrinsic multiplication rate variation of Plasmodium falciparum in clinical isolates prior to elimination in Malaysia" (from another version of this preprint: version 1, 2024-12-12)

**Supplementary Information (2 figures and 3 tables)**

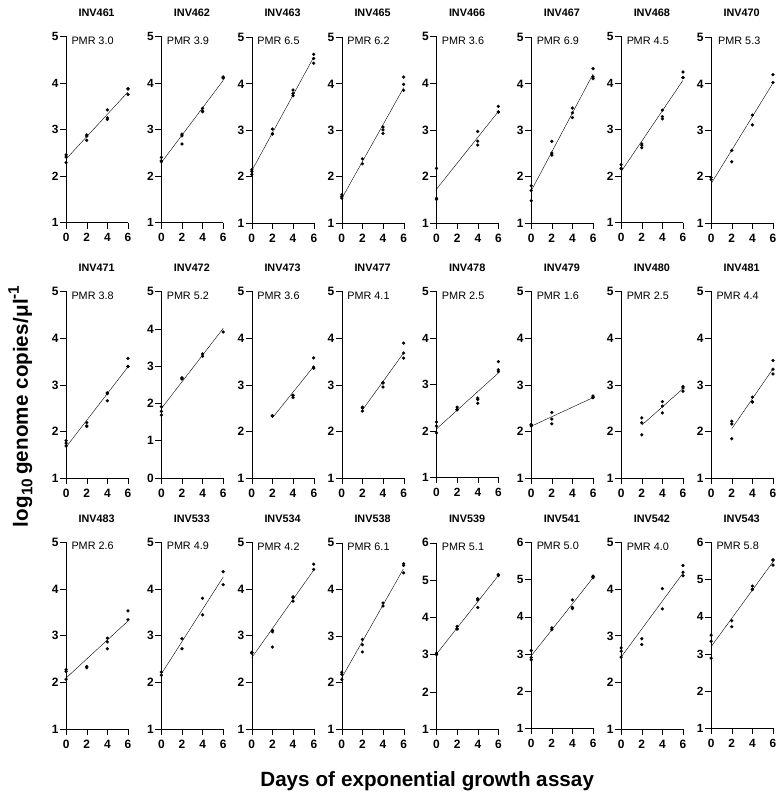

**Supplementary Fig S1.** Exponential multiplication rate assays of 24 Malaysian *P. falciparum* isolates. Each panel shows an individual isolate tested in a 6-day exponential growth assay conducted in triplicate with erythrocytes from three different donors. The plots show parasite genome copies (Log_10_ scale) per microlitre of DNA extracted from each culture timepoint (each 200 microlitre culture sample was extracted into a 50 microlitre volume of DNA). Assays were performed after each isolate had been in continuous culture for least 20 and no more than 23 days, a mean of 21 days (Supplementary Table S1). Parasite multiplication rates (PMR) per 48 hours for each isolate are derived by logistic regression, with 95% confidence intervals of each estimate and other details being shown in Supplementary Table S1. The laboratory-adapted clone 3D7 showed an exponential multiplication rate of approximately 8-fold in the same assays.

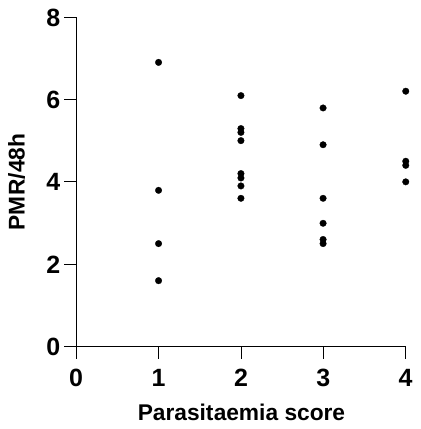

**Supplementary Fig S2.** Correlation of the parasite multiplication rates in culture and the parasitaemia score in patients estimated from thick blood film slide examination in the health facilities (Spearman’s rho = 0.12, P = 0.6, N=22). The patient parasitaemia categories were as follows: 1 = <10 per 100 high power fields (HPF), 2 = 11-100 per 100 HPF, 3 = 11-100 per 10 HPF, 4 = >10 per HPF.

**Supplementary Table S1.** Assayed exponential multiplication rates of *P. falciparum* in cultured isolates from malaria cases in Malaysian Borneo sampled prior to elimination.

| **Isolate Lab ID** | **Local ID** | **Collection site** | **Days in culture at start of 6-day PMR assay** | **Parasite Multiplication Rate per 48 hours (PMR)** | **95% CI of PMR estimate** | **Assay Precision r^2^** |
| --- | --- | --- | --- | --- | --- | --- |
| INV461 | T97/11 | Tawau | 21 | 3.0 | 2.7 - 3.4 | 0.98 |
| INV462 | T97/29 | Tawau | 21 | 3.9 | 3.5 - 4.4 | 0.99 |
| INV463 | T97/07 | Tawau | 21 | 6.5 | 5.9 - 7.2 | 1.00 |
| INV465 | T97/27 | Tawau | 21 | 6.2 | 5.4 - 7.2 | 0.99 |
| INV466 | K96/12 | Kunak | 21 | 3.6 | 2.6 - 5.0 | 0.93 |
| INV467 | K96/10 | Kunak | 21 | 6.9 | 5.8 - 8.1 | 0.98 |
| INV468 | T97/28 | Tawau | 21 | 4.5 | 3.9 - 5.3 | 0.98 |
| INV470 | T97/16 | Tawau | 21 | 5.3 | 4.0 - 7.0 | 0.97 |
| INV471 | L96/04 | Lahad Datu | 21 | 3.8 | 3.3 - 4.3 | 0.98 |
| INV472 | T97/40 | Tawau | 21 | 5.2 | 4.3 - 6.3 | 0.98 |
| INV473 | T97/32 | Tawau | 23 | 3.6 | 2.9 - 4.4 | 0.97 |
| INV477 | T97/10 | Tawau | 23 | 4.1 | 3.3 - 5.2 | 0.97 |
| INV478 | T97/24 | Tawau | 23 | 2.5 | 2.1 - 3.1 | 0.92 |
| INV479 | T97/05 | Tawau | 23 | 1.6 | 1.4 - 1.8 | 0.94 |
| INV480 | T97/36 | Tawau | 23 | 2.5 | 1.9 - 3.2 | 0.90 |
| INV481 | T97/37 | Tawau | 23 | 4.4 | 3.2 - 6.0 | 0.95 |
| INV483 | K96/11 | Kunak | 23 | 2.6 | 2.0 - 3.3 | 0.91 |
| INV533 | T97/03 | Tawau | 20 | 4.9 | 3.8 - 6.5 | 0.97 |
| INV534 | B96/01 | Tawau | 20 | 4.2 | 3.4 - 5.3 | 0.96 |
| INV538 | T97/34 | Tawau | 20 | 6.1 | 5.2 - 7.0 | 0.99 |
| INV539 | T96/10 | Tawau | 20 | 5.1 | 4.6 - 5.6 | 0.99 |
| INV541 | K96/08 | Kunak | 20 | 5.0 | 4.4 - 5.7 | 0.99 |
| INV542 | T97/33 | Tawau | 20 | 4.0 | 3.0 - 5.3 | 0.94 |
| INV543 | L96/16 | Lahad Datu | 20 | 5.8 | 4.5 - 7.4 | 0.97 |

Blood samples of clinical *P. falciparum* infection isolates had been cryopreserved on collection from Kunak and Lahad Datu in 1996, and Tawau in 1997. Culture and assay of the parasites in thawed isolates was performed as described in the Methods after 2021, with triplicate assays of each isolate in different donor erythrocytes. Isolates were processed in three batches (INV461-472, INV473-483, INV533-543), different donor erythrocytes being used in each batch, with no differences seen in the distributions of multiplication rates between batches. The assay data for each isolate are given in Supplementary Table S3.

**Supplementary Table S2 .** Genotypic mixedness of *P. falciparum* isolates in culture as profiled by number of distinct *msp1* and *msp2* alleles detected and parasitaemia scoring of patient clinical samples in the hospitals at time of diagnosis.

|  | **Lab ID** | **Patient parasitaemia score** | | **Mixedness** | **No. of *msp1* alleles** | | **Allelic PCR product sizes**  **(bp)** | | | **No. of *msp2* alleles** | | **Allelic PCR product sizes**  **(bp)** | | | | | **MOI** |
| --- | --- | --- | --- | --- | --- | --- | --- | --- | --- | --- | --- | --- | --- | --- | --- | --- | --- |
|  | INV461 | 3+ | | MIXED | 1 | | R-156 | | | 2 | | IC-447,531 | | | | | 2 |
|  | INV462 | 2+ | | MIXED | 1 | | R-156 | | | 2 | | IC-459,540 | | | | | 2 |
|  | INV463 | - | | SINGLE | 1 | | M-234 | | | 1 | | FC - 327 | | | | | 1 |
|  | INV465 | 4+ | | SINGLE | 1 | | M-206 | | | 1 | | FC - 327 | | | | | 1 |
|  | INV466 | 3+ | | MIXED | 2 | | M-206,240 | | | 2 | | IC-465,537 | | | | | 2 |
|  | INV467 | 1+ | | SINGLE | 1 | | R-156 | | | 1 | | FC-327 | | | | | 1 |
|  | INV468 | 4+ | | MIXED | 2 | | M-159,R156 | | | 1 | | FC-327 | | | | | 2 |
|  | INV470 | 2+ | | SINGLE | 1 | | M-240 | | | 1 | | FC-333 | | | | | 1 |
|  | INV471 | 1+ | | SINGLE | 1 | | M-213 | | | 1 | | FC-333 | | | | | 1 |
|  | INV472 | 2+ | | SINGLE | 1 | | M-216 | | | 1 | | IC-666 | | | | | 1 |
|  | INV473 | 2+ | | MIXED | 1 | | R-156 | | | 2 | | IC-432,501 | | | | | 2 |
|  | INV477 | 2+ | | SINGLE | 1 | | M-168 | | | 1 | | FC-333 | | | | | 1 |
|  | INV478 | 3+ | | SINGLE | 1 | | K-192 | | | 1 | | FC-312 | | | | | 1 |
|  | INV479 | 1+ | | SINGLE | 1 | | R-156 | | | 1 | | FC-315 | | | | | 1 |
|  | INV480 | 1+ | | SINGLE | 1 | | M-162 | | | 1 | | FC-333 | | | | | 1 |
|  | INV481 | 4+ | | MIXED | 1 | | R-156 | | | 3 | | FC-333,456,513 | | | | | 3 |
|  | INV483 | 3+ | | MIXED | 2 | | M-186,R-156 | | | 2 | | IC-465,531 | | | | | 2 |
|  | INV533 | 3+ | | SINGLE | 1 | | R-156 | | | 1 | | IC-636 | | | | | 1 |
|  | INV534 | 2+ | | SINGLE | 1 | | M-195 | | | 1 | | FC- 288 | | | | | 1 |
|  | INV538 | 2+ | | MIXED | 2 | | M-210,R-156 | | | 1 | | FC-300 | | | | | 2 |
|  | INV539 | - | | SINGLE | 1 | | R-156 | | | 1 | | FC-300 | | | | | 1 |
|  | INV541 | 2+ | | SINGLE | 1 | | M-219 | | | 1 | | FC-309 | | | | | 1 |
|  | INV542 | 4+ | | SINGLE | 1 | | M-240 | | | 1 | | IC-669 | | | | | 1 |
|  | INV543 | 3+ | | SINGLE | 1 | | M-225 | | | 1 | | IC-324 | | | | | 1 |
| Genotyping was performed on DNA extracted from cultures at or shortly before the time of the multiplication rate assays, by nested PCR as described in the Methods.  Patient parasitaemia categorical scores estimated by microscopy at time of diagnosis in health facilities:  1+, <10 per 100 high power fields (HPF); 2+, 11-100 per 100 HPF; 3+, 11-100 per 10 HPF; 4+, >10 per HPF. | | | | | | | | | | | | | | | | | |

**Supplementary Table S3.** Quantitative PCR assay data on genome copies per microlitre of DNA extracted from each of the triplicate assay points for each of the isolates in the exponential growth rate assays.

|  | Isolate ID (INV) numbers are given in the headings and genome copies on each assay day in vertical columns for the different donor erythrocytes. Any missing data points show as dashes (-) | | | | | | | | | | | |
| --- | --- | --- | --- | --- | --- | --- | --- | --- | --- | --- | --- | --- |
| **Days** | **461** | | | **462** | | | **463** | | | **465** | | |
| **0** | 776 | 1138 | 1030 | 1011 | 812 | 850 | 518 | 564 | 446 | 139 | 163 | 150 |
| **2** | 3036 | 2883 | 2355 | 3196 | 2927 | 1956 | 3295 | 4200 | 3359 | 758 | 967 | 766 |
| **4** | 10620 | 7141 | 6582 | 9948 | 9693 | 11500 | 21960 | 24460 | 29060 | 3389 | 4687 | 4062 |
| **6** | 29820 | 30260 | 22670 | 54130 | 51360 | 51510 | 109500 | 138200 | 170700 | 38450 | 55370 | 28650 |
| **Days** | **466** | | | **467** | | | **468** | | | **470** | | |
| **0** | 597 | 129 | 136 | 256 | 202 | 122 | 588 | 704 | 577 | 386 | 347 | - |
| **2** | - | - | - | 2299 | 1165 | 1292 | 1850 | 1970 | 1634 | 837 | 1456 | - |
| **4** | 1877 | 3697 | 2288 | 11900 | 9421 | 7453 | 10500 | 7629 | 6785 | 5175 | 8376 | - |
| **6** | 9662 | 12700 | 9596 | 83800 | 56000 | 51100 | 52600 | 69500 | 52600 | 42400 | 62200 | - |
| **Days** | **471** | | | **472** | | | **473** | | | **477** | | |
| **0** | 194 | 223 | 252 | 326 | 243 | 193 | - | - | - | - | - | - |
| **2** | 614 | 523 | 509 | 1813 | 1932 | 1912 | 870 | 855 | 850 | 1275 | 1319 | 1086 |
| **4** | 2695 | 2574 | 1807 | 8449 | - | 7392 | 2111 | 2379 | 2344 | 4402 | 3577 | 4359 |
| **6** | 9844 | 14700 | 9805 | - | 32900 | - | 9581 | 9015 | 15100 | 14900 | 19100 | 31100 |
| **Days** | **478** | | | **479** | | | **480** | | | **481** | | |
| **0** | 509 | 364 | 623 | 518 | 563 | 548 | - | - | - | - | - | - |
| **2** | 1303 | 1141 | 1147 | 732 | 1012 | 582 | 779 | 612 | 336 | 278 | 657 | 578 |
| **4** | 1904 | 1574 | 2030 | - | - | - | 1726 | 1389 | 995 | 1723 | 2177 | 1707 |
| **6** | 8289 | 7586 | 12400 | 2100 | 2158 | 2285 | 3429 | 3684 | 2890 | 6838 | 13200 | 8632 |
| **Days** | **483** | | | **533** | | | **534** | | | **538** | | |
| **0** | 683 | 461 | 750 | 1753 | 1802 | 2373 | 1713 | 1755 | 1694 | 663 | 462 | 601 |
| **2** | 871 | 820 | - | - | 6908 | 4390 | 2265 | 5160 | 4833 | 1806 | 3347 | 2585 |
| **4** | 2923 | 3538 | 2084 | 45400 | 62300 | 122000 | 21900 | 27200 | 26100 | 20390 | - | 17600 |
| **6** | 8764 | - | 13600 | 319000 | - | 145000 |  | 105000 | 136000 | 90600 | 142000 | 130000 |
| **Days** | **539** | | | **541** | | | **542** | | | **543** | | |
| **0** | 4084 | 4318 | 4034 | 3203 | 4958 | 2794 | 1391 | 1861 | 2189 | 3091 | 8798 | 12770 |
| **2** | 19100 | 22900 | 19700 | - | 20100 | 18000 | - | 2583 | 3447 | - | 21630 | 31230 |
| **4** | 126000 | 73100 | 117000 | 71400 | 113000 | 66300 | 41000 | - | 15100 | 217600 | 214200 | 265600 |
| **6** | 561000 | 528000 | - | 478000 | 492000 | 447000 | 129000 | 91500 | 77600 | 1335000 | 979000 | 1353000 |
